# A three-component logic gate governs quorum sensing-regulated killing of *Stenotrophomonas maltophilia* by *Pseudomonas aeruginosa*

**DOI:** 10.64898/2026.01.27.702039

**Authors:** Andrew Frando, Robert S. Parsek, Graham W. Roberts, Ajai A. Dandekar

**Author notes:** Address correspondence to: Ajai A. Dandekar, K-359A HSB, Box 356522, 705 NE Pacific St Seattle, WA 98195.

## Abstract

*Pseudomonas aeruginosa*, an opportunistic pathogen, uses a trio of quorum sensing (QS) systems to regulate the production of some virulence factors. Two of these, the *las* and *rhl* systems, involve acyl-homoserine lactone signals; the third, called *pqs*, primarily uses the signal 2-heptyl-3-hydroxy-4(1*H*)-quinolone (“PQS”). We aimed to identify how interbacterial interactions are regulated between *P. aeruginosa* and *Stenotrophomonas maltophilia*, which co-occur in many environments, including the airways of people with cystic fibrosis. We explored *P. aeruginosa* and *S. maltophilia* interactions using a co-culture model. In the conditions of our experiments, *P. aeruginosa* kills *S. maltophilia*. Co-culture of *S. maltophilia* with *P. aeruginosa* deficient in *las*, *rhl*, or *pqs* QS resulted in greater *S. maltophilia* viability than co-culture with the wildtype. This inhibition was not generally attributable to *las* and *rhl*-regulated toxins. Therefore, we interrogated the role of *pqs* QS and found that co-culture of *S. maltophilia* with *P. aeruginosa* deficient in PQS biosynthesis showed similar CFUs to monoculture. Exogenous PQS did not complement this phenotype, suggesting that another quinolone is the effector. We found that *S. maltophilia* killing is reduced in competition with a mutant that cannot make the quinolone HQNO (2-heptyl-4-quinoline *N*-oxide). We show that full killing of *S. maltophilia* by *P. aeruginosa* requires three components: HQNO, the chaperone PqsE, and intact PQS biosynthesis that together form a tripartite AND logic gate. Our work identifies quinolone quorum sensing as a driver for interactions between the Gram-negative pathogens *P. aeruginosa* and *S. maltophilia*.

## Introduction

Bacteria are typically found in microbial communities where they interact with biotic and abiotic factors, as in the soil and gastrointestinal tract of animals [1, 2]. Bacteria can interact with other microbes in these spaces, and these interactions can alter their behaviors [3–6]. These behaviors can affect their immediate environment, including nearby bacteria. The Gram-negative bacterium *Pseudomonas aeruginosa* occupies a variety of environmental niches that it can share with other bacteria, including the soil, hospital sink drains, and animal hosts [7–10]. *P. aeruginosa* can be co-isolated from human infections with other bacteria, including *Staphylococcus aureus* and *Stenotrophomonas maltophilia* [11–15]. In this situation, *P. aeruginosa* produces several secreted factors, including those involved in virulence, that have been shown to affect human hosts and its interactions with other microbes [16–21].

Interactions between *P. aeruginosa* and some other bacteria, such as the Gram-positive bacterium *S. aureus*, have been well-described. For example, *P. aeruginosa* in coculture with *S. aureus* secretes virulence factors including the alkylquinolone 2-heptyl-4-quinoline *N*-oxide (HQNO) and the siderophores pyoverdine and pyochelin [22]. These virulence factors negatively impact *S. aureus* metabolism, allowing *P. aeruginosa* to gain a competitive advantage and increase its growth [23, 24]. While this interaction seems to be more beneficial for *P. aeruginosa*, growth together results in *S. aureus* having increased resistance to antibiotics, including gentamicin and tetracycline [25, 26]. This finding parallels other studies which show that interactions between *P. aeruginosa* and other microbes can increase *P. aeruginosa* growth [24, 27], result in increased drug resistance [25, 26, 28], or both; all of these outcomes likely are detrimental to infected hosts. These studies highlight the importance of understanding interbacterial interactions and how they affect bacterial behaviors and phenotypes.

Interactions between *P. aeruginosa* and Gram-negative bacteria have been less well-characterized, although these bacteria frequently co-occur [7–10]. One such bacterium that can share niches with *P. aeruginosa* is the saprophyte *Stenotrophomonas maltophilia.* Interactions between *P. aeruginosa* and *S. maltophilia* have been sparsely studied, mostly in the context of human infections of people with the genetic disease cystic fibrosis (CF). *S. maltophilia* is co-isolated with *P. aeruginosa* at a frequency ranging from 10 to 60% in CF airways [29–32]. Interactions between *P. aeruginosa* and *S. maltophilia* may influence the antimicrobial resistance profile of *P. aeruginosa* [33].

*P. aeruginosa* regulates behaviors within the species using an intercellular signaling system known as quorum sensing (QS). Some bacteria use QS to assess their cell density and to alter population behaviors in response [34]. In *P. aeruginosa*, QS regulates the production of secreted products like the protease elastase, toxic biosurfactant rhamnolipids, and hydrogen cyanide [35–38]. QS also partially regulates the production of extracellular polysaccharides that are important for forming biofilms [39, 40]. In general, QS consists of a circuit composed of a secreted signal; this signal is sensed by and activates a regulator. *P. aeruginosa* harbors three complete QS circuits that involve the transcription factors LasR, RhlR, and PqsR (also called MvfR) [41–43]. These circuits use different signals. LasR binds to the signal *N*-3-oxo-dodecanoyl-homoserine lactone (3OC12-HSL) produced by the signal synthase LasI; similarly, RhlR binds to *N-*butanoyl-homoserine lactone (C4-HSL) produced by RhlI. PqsR recognizes 2-heptyl-3-hydroxy-4(1*H*)-quinolone (called the Pseudomonas quinolone signal, PQS) which is synthesized by the products of an operon, *pqsABCDE* and an unlinked gene, *pqsH* [44–46] (**Supplemental Figure 1**). These three QS circuits (*las*, *rhl*, and *pqs)* control the expression of hundreds of genes, including several that affect virulence, and they are arranged in a hierarchy: LasR activates expression of the genes encoding RhlR and PqsR [35, 43, 47, 48].

*las* and *rhl* QS are known to regulate interactions with Gram-negative bacteria [19–21], but the role of *pqs* QS in these interactions is less well-described. The enzymes of PQS biosynthesis together modify chorismic acid into PQS. Each enzyme controls the production of an intermediate in this pathway, and mutants in any gene in the PQS biosynthesis pathway (except PqsE) are unable to produce PQS (**Supplemental Figure 1**) [49]. Signal-bound PqsR positively regulates the expression of *pqsABCDE* and thus creates a positive feedback loop. PqsE, a product of this operon, is unique: it is required to fully activate *rhl* QS [50], and in many strains is dispensable for PQS biosynthesis [45, 46, 51]. Finally, the enzyme PqsL, a product of another gene unlinked to *pqsABCDE*, creates a branch in the PQS biosynthesis pathway and is required to produce HQNO (**Supplemental Figure 1**). PQS biosynthesis is unaffected in a *pqsL* mutant [46, 52]. HQNO and other quinolones (there are dozens produced as intermediates in this pathway) have been demonstrated to have antibiotic activity against a range of bacteria, including *S. aureus* [16, 22, 26, 53, 54].

We are interested in understanding the interactions of *P. aeruginosa* and *S. maltophilia*, to better define the determinants of competition between Gram-negative bacteria, and the genes that modulate these interactions. We seek to understand how these interactions affect the surrounding environment and microbes. We established a co-culture system and determined that, in the conditions of our experiments, *P. aeruginosa* kills *S. maltophilia*. We assessed the genes affecting this interaction and found that inactivation of any of the three QS circuits abrogated the killing of *S. maltophilia*. Because *pqs* QS is regulated by LasR, and its contributions to interactions with Gram-negative bacteria are relatively poorly understood, we explored its role further. We found that the inhibitor HQNO, the chaperone PqsE, and PQS biosynthesis are all necessary for mediating competition between *P. aeruginosa* and *S. maltophilia*. These data have implications for how bacteria use alkylquinolones to mediate interactions with their environments, the integration of multiple cellular pathways to coordinate interactions with other bacteria, and generally how Gram-negative bacteria interact in a variety of environments, including during infection and in natural settings.

## Results

### P. aeruginosa competes with S. maltophilia

We developed a co-culture system using a condition where the *P. aeruginosa* strain PAO1 and the *S. maltophilia* strain K279a grew in monoculture with similar growth rates and yields (**Supplemental Figure 2**). We monitored bacterial viability using colony forming unit (CFU) assays on selective plates: we selected for *S. maltophilia* using LB agar supplemented with gentamicin and selected for *P. aeruginosa* using Columbia agar supplemented with C-390 and phenanthroline [55]. When these bacteria were grown in co-culture, we found that *S. maltophilia* viability was unchanged as compared to monoculture until approximately 12 hours, where we observed a ∼1-log decrease (**Figure 1A**). At 16 hours, we observed a sharper decline in CFUs. By 20 hours, we observed a striking phenotype where *S. maltophilia* viability decreased by ∼7-logs in co-culture with *P. aeruginosa* compared to monoculture. *S. maltophilia* viability remained decreased until the end of the experiment. We found that *P. aeruginosa* viability was unchanged in the same co-culture conditions compared to monoculture (**Figure 1B**). These findings demonstrated that *S. maltophilia* and *P. aeruginosa* interact and that *P. aeruginosa* kills *S. maltophilia* by some mechanism.

**Figure 1.**
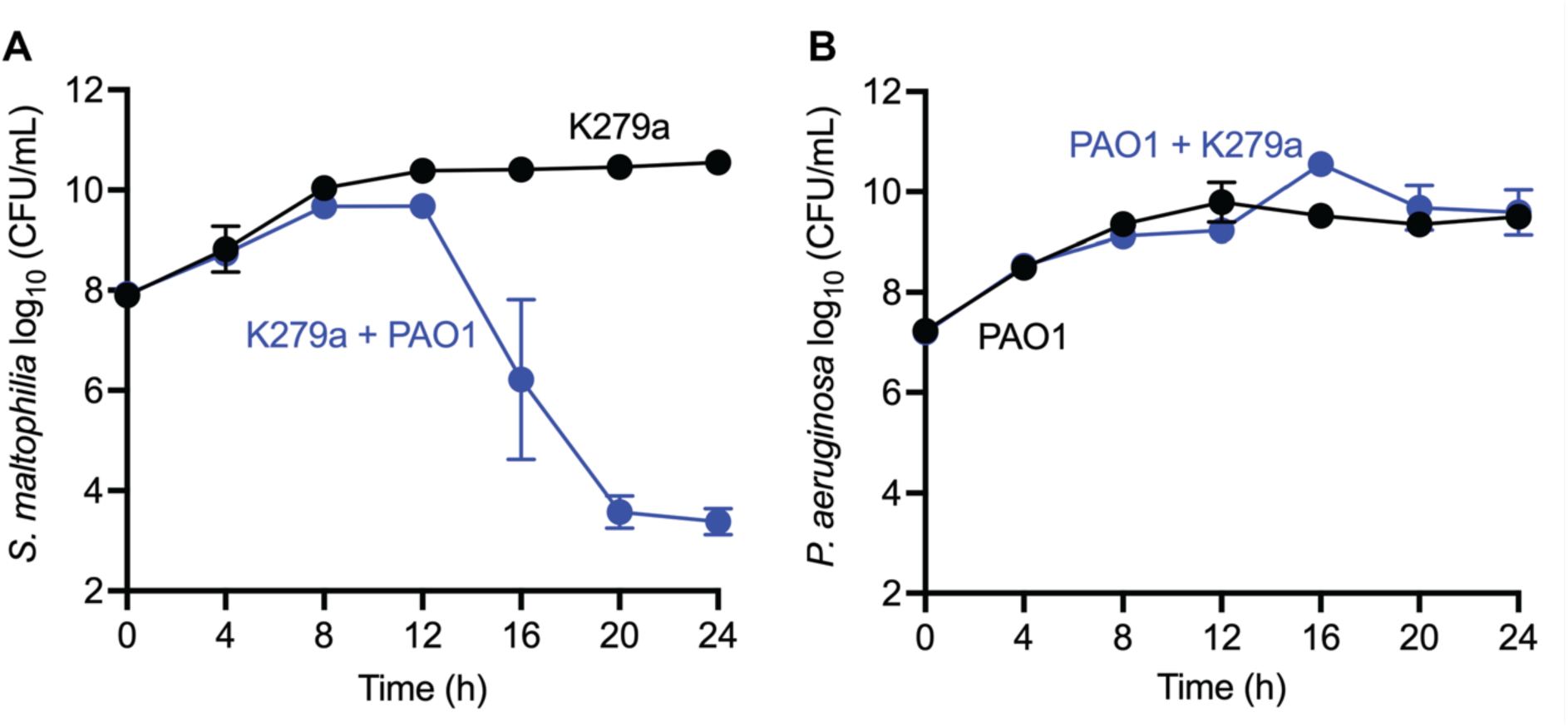
*P. aeruginosa* kills *S. maltophilia* in co-culture while *P. aeruginosa* viability is mostly unaffected. Wild-type *P. aeruginosa* PAO1 and wild-type *S. maltophilia* K279a were grown in mono- or co-culture for 24 h. In co-cultures, PAO1 and K279a were mixed at a 1:1 ratio based on OD_600_. *S. maltophilia* CFUs are shown in **A** and *P. aeruginosa* CFUs are shown in **B**. The experiment was performed in triplicate. Error bars represent the standard deviation. *P* values were calculated between mono- and co-cultures using a two-way ANOVA with Geisser-Greenhouse correction. The curves in panel A were statistically different (*P* < 0.0001), those in panel B were not (*P* = 0.1486).

### Competition between *P. aeruginosa* and *S. maltophilia* is mediated by quorum sensing

We were next asked what genes might be mediating interactions between *P. aeruginosa* and *S. maltophilia*. Competition between *P. aeruginosa* and some other bacteria has been demonstrated to be regulated by QS, so we asked if QS played a role in this competition. To answer this question, we grew *S. maltophilia* K279a in mono- or co-culture with wild-type PAO1, PAO1Δ*lasR,* PAO1Δ*rhlR,* or PAO1Δ*pqsR* and measured *S. maltophilia* CFUs after 24 hours **(Figure 2)**. In contrast to co-culture with wild-type *P. aeruginosa*, we found that *S. maltophilia* exhibited markedly increased viability when co-cultured with mutants in each of the QS transcriptional regulators. *S. maltophilia* co-cultured with either the *lasR* or *rhlR* mutants had a 5-log increase and the *pqsR* mutant had a 6-log increase in *S. maltophilia* viability, as compared to the wild-type. These data show that *S. maltophilia* is better able to compete against *P. aeruginosa* QS mutants, demonstrating that *P. aeruginosa* QS mediates competition with *S. maltophilia*.

**Figure 2.**
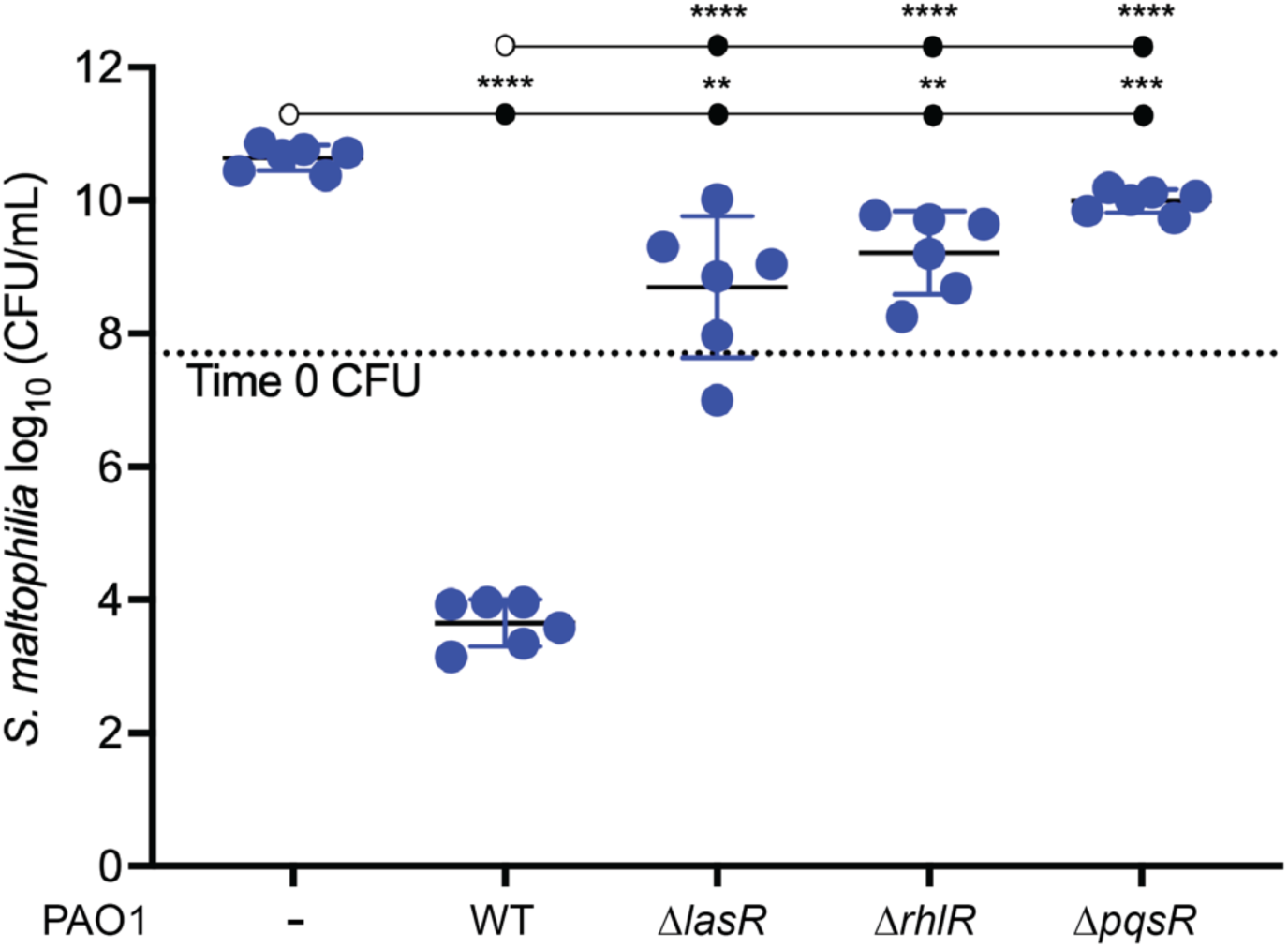
Competition between *P. aeruginosa* and *S. maltophilia* is mediated by quorum sensing. Wild-type *S. maltophilia* K279a was grown in mono- or co-culture with wild-type *P. aeruginosa* PAO1, PAO1Δ*lasR*, PAO1Δ*lasR*, PAO1Δ*rhlR*, or PAO1Δ*pqsR*. The graph shows *S. maltophilia* viability at 24 hours where the x-axis shows each condition and the y-axis shows *S. maltophilia* CFUs. The experiment was performed in triplicate. Error bars represent the standard deviation. *P* values were calculated between each strain using an unpaired two-tailed t test with Welch’s correction. Statistical comparisons between strains are indicated by a line connecting open and closed circles, with ** denoting *P* < 0.01, *** denoting *P* < 0.001, and **** denoting *P* < 0.0001.

Prior work has shown that several *P. aeruginosa* LasR- and RhlR-regulated factors are important for mediating competition with other bacteria. These products include phenazines such as pyocyanin, hydrogen cyanide, and the biosurfactants rhamnolipids. We tested whether these QS-regulated factors were responsible for the competitive phenotype by co-culturing *S. maltophilia* with mutants deficient in phenazine biosynthesis (PAO1Δ*phzA1*), hydrogen cyanide production (PAO1Δ*hcnC*), or rhamnolipid biosynthesis (PAO1Δ*rhlB*) (**Supplemental Figure 3**). We found that *P. aeruginosa* mutants unable to produce hydrogen cyanide or rhamnolipids had only minor competitive defects when co-cultured with *S. maltophilia*. Further, we found that *P. aeruginosa* mutants unable to produce phenazines exhibited similar competition with *S. maltophilia* as the wild-type. We concluded that while these factors might explain a small component of the competition with *S. maltophilia*, they do not account for large competitive defects we observed with the QS transcriptional regulator mutants.

### Alkylquinolones by themselves do not mediate competition with *S. maltophilia*

LasR activates *pqsR* expression, and therefore a *lasR* mutant does not exhibit PqsR activity [44, 56]. Having excluded major LasR- and RhlR-regulated factors as mediators of *S. maltophilia* killing, we reasoned that the competitive defect in the *lasR* mutant (**Figure 2**) may be due to a resultant lack of *pqs* QS. Alkylquinolones that are produced as part of *pqs* QS and PQS biosynthesis have been shown to mediate interbacterial interactions, such as HQNO-mediated growth inhibition of *S. aureus* [16, 22, 26]. Therefore, we explored the role of *pqs* QS in mediating interactions between *P. aeruginosa* and *S. maltophilia*.

We first asked whether alkylquinolones could mediate competition with *S. maltophilia*, as in the case of *S. aureus*. Some alkylquinolones are commercially available, including the signaling molecules PQS and HHQ and the inhibitor HQNO; therefore we treated *S. maltophilia* monocultures with purified PQS, HHQ, or HQNO at concentrations previously determined in stationary phase cultures [46, 53, 57] (**Figure 3A**). We found no change in *S. maltophilia* viability when treated with each of the alkylquinolones. Since *P. aeruginosa* produces all three of the alkylquinolones during growth, we next probed whether the alkylquinolones mediate competition when added in combination. We discovered that even when added in dual or triple combinations, *S. maltophilia* viability was unchanged compared to the untreated control. These data indicate that alkylquinolones alone or in combination are insufficent to effect killing of *S. maltophilia* by *P. aeruginosa*.

**Figure 3.**
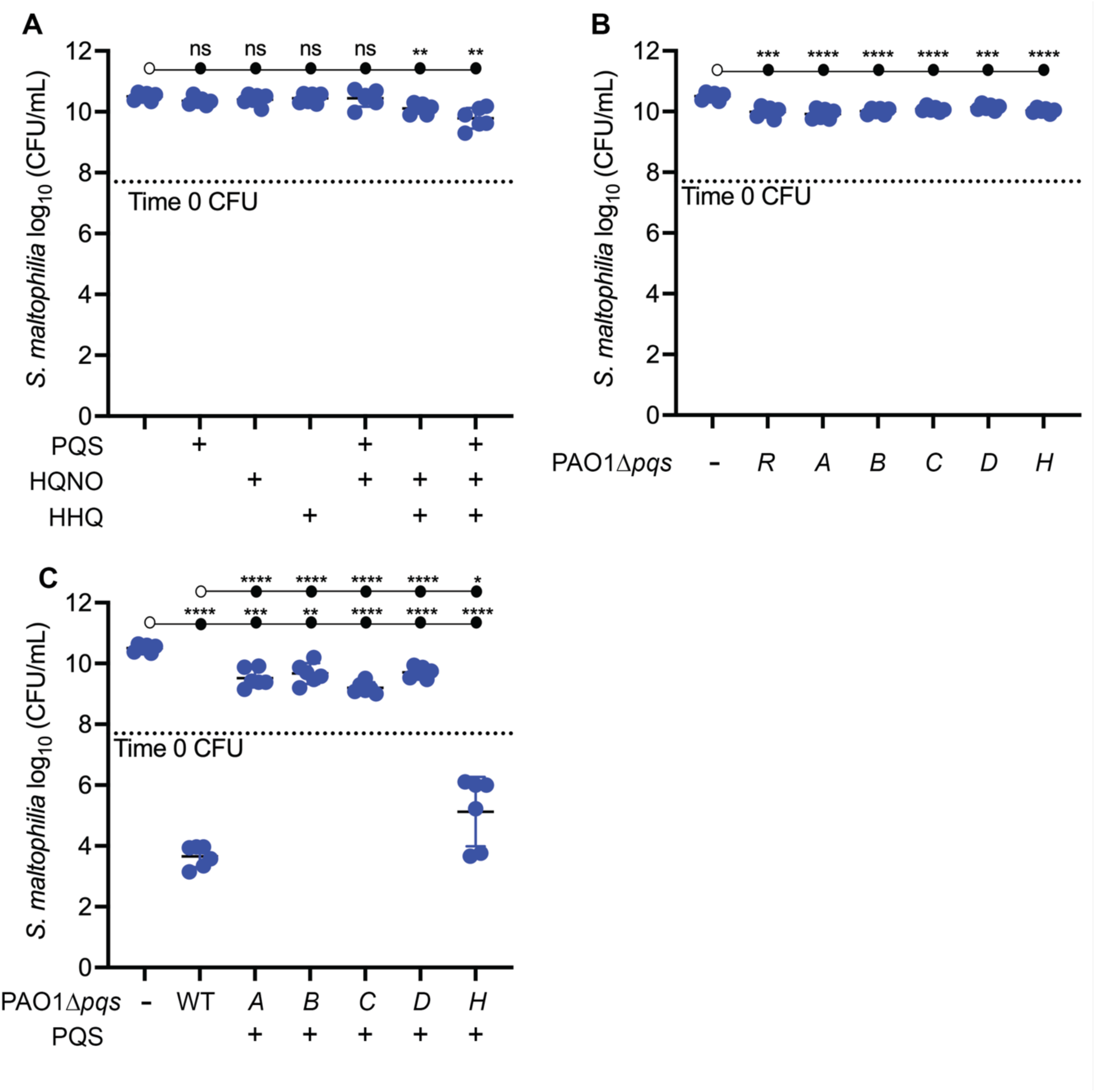
*pqs* QS, but not alkylquinolones alone, mediates competition between *P. aeruginosa* and *S. maltophilia*. **A**. Wild-type K279a cultures were supplemented with 20 µM 2-heptyl-3-hydroxy-4(1H)-quinolone (PQS), 50 µM 2-heptyl-4-quinolone (HHQ), or 50 µM 2-heptyl-4-hydroxyquinoline *N*-oxide (HQNO) individually or in combination. Cultures were grown for 24 hours. **B**. Wild-type K279a was grown in mono- or co-culture with wild-type PAO1, PAO1Δ*pqsR*, PAO1Δ*pqsA*, PAO1Δ*pqsB*, PAO1Δ*pqsC*, PAO1Δ*pqsD*, or PAO1Δ*pqsH*. **C**. Wild-type K279a was grown in mono- or co-culture with wild-type PAO1, PAO1Δ*pqsR*, PAO1Δ*pqsA*, PAO1Δ*pqsB*, PAO1Δ*pqsC*, PAO1Δ*pqsD*, or PAO1Δ*pqsH* supplemented with 20 µM PQS. All graphs show *S. maltophilia* viability at 24 hours where the x-axis shows each condition and the y-axis shows *S. maltophilia* CFUs. Each experiment was performed in triplicate. Error bars represent the standard deviation. *P* values were calculated between each condition using an unpaired two-tailed t test with Welch’s correction. Statistical comparisons between strains are indicated by a line connecting open and closed circles, with * denoting *P* < 0.05, ** denoting *P* < 0.01, *** denoting *P* < 0.001, and **** denoting *P* < 0.0001.

### Products of *pqs* QS mediate competition between *P. aeruginosa* and *S. maltophilia*

The *pqs* QS pathway regulates the expression of the PQS biosynthesis locus (*pqsABCDE and phnAB)* that not only produces the compounds PQS, HHQ, HQNO, but also over 20 intermediates required to produce these compounds [46, 58]. We next wanted to probe the role of these compounds and test the role of the PQS biosynthesis locus in mediating competition between *P. aeruginosa* and *S. maltophilia*. To do so, we took a genetic approach by generating deletion mutants in each of the PQS biosynthesis genes and competed these mutants against *S. maltophilia* (**Figure 3B**). Identical to PAO1Δ*pqsR*, we observed that mutants of each of the PQS biosynthesis genes had a large competitive defect when grown with *S. maltophilia*, whose growth and viability was only slightly less than in monoculture. These data are consistent with the idea that *pqs* QS, specifically a product or products of the PQS biosynthesis locus, mediates competition between *P. aeruginosa* and *S. maltophilia*.

We next wanted to identify if any specific genes in the PQS biosynthesis locus, and possibly their products, mediated competition. Gene products of the PQS biosynthesis locus produce a series of intermediates that ends in PQS. PQS is the major quinolone that binds to and activates PqsR, which in turn upregulates expression of the PQS biosynthesis locus, creating a positive feedback loop. Therefore, a mutant in any of the PQS biosynthesis locus genes disrupts the production of PQS, the intermediate alkylquinolone products, and also the positive feedback loop required for QS. To test the role of the specific gene in the PQS biosynthesis locus while also having the feedback loop remain active, we competed individual PQS biosynthesis operon mutants against *S. maltophilia* and added PQS at the beginning of the experiment (**Figure 3C**). Adding PQS relieves the defect in *pqsABCDE* expression but not the production of alkylquinolones in these mutants. We observed that addition of PQS to *pqsA*, *pqsB*, *pqsC*, and *pqsD* mutants did not change the competition phenotype, indicating that these genes and their products were insufficient to kill *S. maltophilia*. Interestingly, the *pqsH* mutant (which is only defective in production of PQS itself) supplemented with exogenous PQS regained the ability to compete with *S. maltophilia*; we observed a ∼4-log decrease in *S. maltophilia* CFUs when PQS was added compared to when PQS was absent. These data indicate that a product of *pqs* QS was responsible for competition between *P. aeruginosa* and *S. maltophilia*, although this product may not be among the enzymes that synthesize intermediates required for producing PQS.

### HQNO is required for competition between *P. aeruginosa* and *S. maltophilia*

HQNO is another terminal product of alkylquinolone synthesis that is produced by the enzyme PqsL (**Supplemental Figure 1**). This is a branch from the PQS biosynthesis pathway. PqsL diverts some of the 2-ABA away from PQS biosynthesis by converting of 2-ABA to a hydroxylamino derivative of 2-ABA that is subsequently converted to HQNO by the enzymes PqsB and PqsC. As such, we assessed the role of PqsL and its products in mediating competition between *P. aeruginosa* and *S. maltophilia* by growing *S. maltophilia* in co-culture with a PAO1Δ*pqsL* mutant (**Figure 4A**). The *pqsL* mutant is only deficient in its ability to produce HQNO but it can still synthesize PQS and intermediate alkylquinolone products and activate *pqs* QS. This *pqsL* mutant, unlike a *pqsR* deletion mutant, still killed *S. maltophilia*, although not as well as the WT (**Figure 4A**). Compared to *S. maltophilia* monoculture, we observed a ∼3-log decrease in *S. maltophilia* CFU when co-cultured with the *pqsL* deletion mutant.

**Figure 4.**
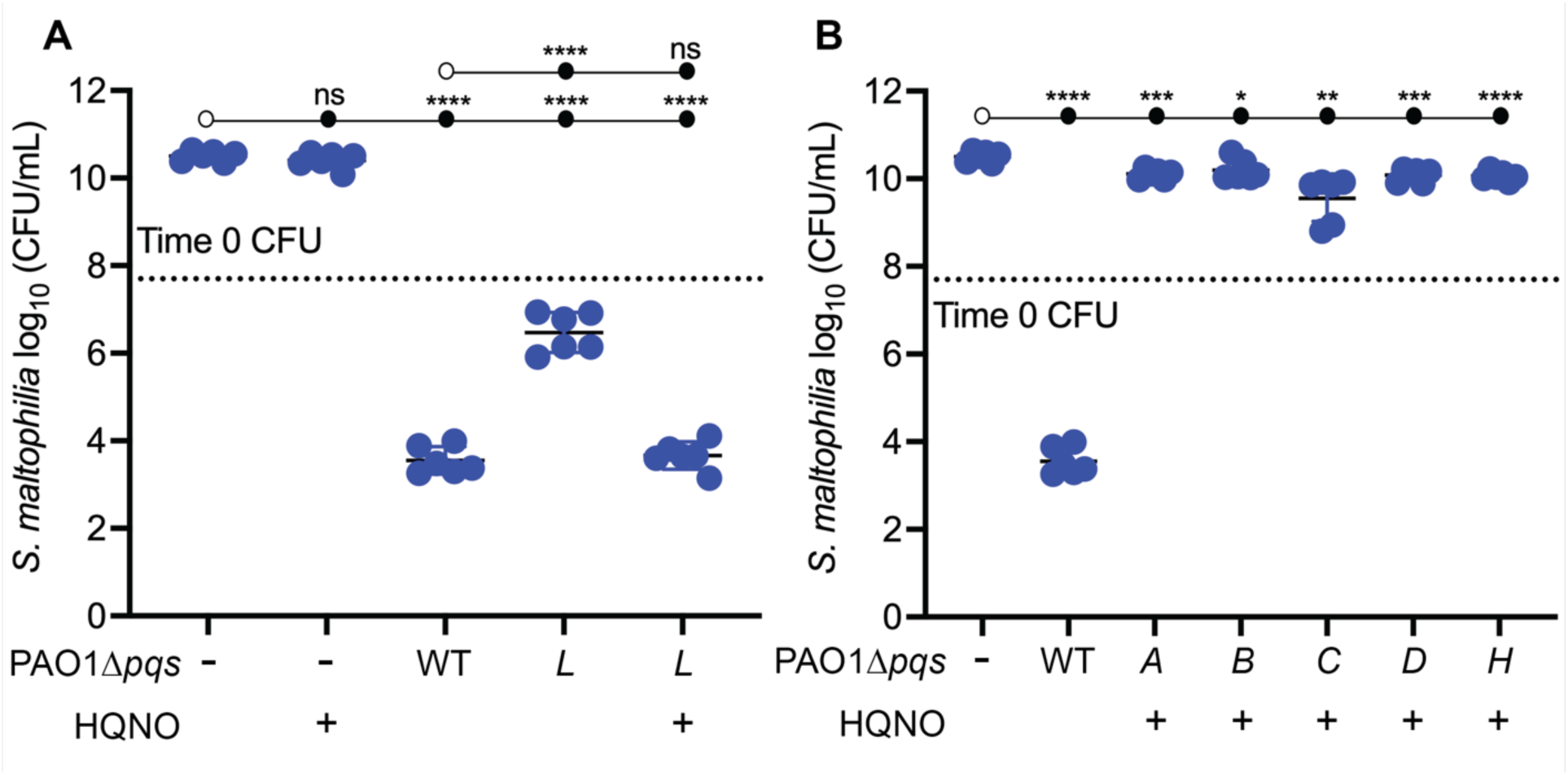
Competition between *P. aeruginosa* and *S. maltophilia* is partially mediated by HQNO and requires *pqs* QS. **A.** Wild-type K279a was grown in mono- or co-culture with wild-type PAO1 or PAO1Δ*pqsL* supplemented with 50 µM HQNO. **B.** Wild-type K279a was grown in mono- or co-culture with wild-type PAO1, PAO1Δ*pqsA*, PAO1Δ*pqsB*, PAO1Δ*pqsD*, or PAO1Δ*pqsH* supplemented with 50 µM HQNO. All graphs show *S. maltophilia* viability at 24 hours where the x-axis shows each condition and the y-axis shows *S. maltophilia* CFUs. Each experiment was performed in triplicate. Error bars represent the standard deviation. *P* values were calculated between each condition using an unpaired two-tailed t test with Welch’s correction. Statistical comparisons between strains are indicated by a line connecting open and closed circles, with * denoting *P* < 0.05, ** denoting *P* < 0.01, *** denoting *P* < 0.001, and **** denoting *P* < 0.0001.

Because this *pqsL* mutant is only deficient in HQNO production, the increase in *S. maltophilia* viability as compared to the WT was likely attributable to HQNO. To test this idea, we added exogenous HQNO to the *S. maltophilia* - PAO1Δ*pqsL* co-culture. We observed that, with the addition of HQNO, *S. maltophilia* viability was identical to that seen in co-culture with WT *P. aeruginosa*. This result indicated that HQNO can complement the competitive defect seen in the *pqsL* mutant and that HQNO mediates an element of the competition between *P. aeruginosa* and *S. maltophilia*.

The result that HQNO is important for *S. maltophilia* killing was seemingly contrary to our prior data: when we added HQNO to *S. maltophilia* monoculture, we did not observe any changes in viability (**Figure 3A**), unlike what has been shown for the Gram-positive bacterium *S. aureus [16, 22, 26]*. To investigate this paradox, we asked whether *S. maltophilia* killing by any of the other mutants of the PQS biosynthesis locus could be complemented with the addition of HQNO (**Figure 4B**). We found that the addition of HQNO did not change *S. maltophilia* viability in co-culture with any of these mutants. Because the *pqsL* mutant is the only mutant in the PQS and HQNO biosynthesis pathway that maintains the ability to activate *pqs* QS, we reasoned that it was likely that killing by HQNO is dependent on the presence of another *pqs* QS-regulated factor.

### A PqsE-regulated factor potentiates HQNO activity

We therefore aimed to pinpoint the identity of this other factor that could potentiate HQNO activity. Our data thus far indicated that *pqs* QS was important, in addition to HQNO activity, for killing (**Figure 4**). We turned our attention to PqsE. PqsE has thioesterase activity and can catalyze the conversion of 2-aminobenzoylacetyl-coenzyme A to 2-aminobenzoylacetate; however, it is disposable for PQS biosynthesis in PAO1 [51, 59]. PqsE also links the *pqs* and *rhl* QS pathways by serving a chaperone-like function for RhlR [60, 61]; there are RhlR-regulated genes (such as those encoding for phenazine biosynthesis) whose expression are exquisitely dependent on PqsE [62–65].

We assessed the individual role of PqsE and, by connection, RhlR, activity in modulating competition between *P. aeruginosa* and *S. maltophilia* (**Figure 5A**). The *pqsE* mutant killed *S. maltophilia*, but not as well as the WT, resulting in a ∼3-log decrease in *S. maltophilia* CFU as compared to monoculture growth. We next tested whether the *pqsE* mutant could be complemented with the addition of exogenous HQNO (**Figure 5A**). We did not observe any changes in *S. maltophilia* viability when HQNO was added to the co-culture with PAO1Δ*pqsE*, which was not a surprising result, as a *pqsE* mutant can still synthesize HQNO (**Supplemental Figure 1**). These results collectively indicated a role for PqsE in competition and suggest that factors regulated by PqsE may be important for potentiating HQNO activity against *S. maltophilia*.

**Figure 5.**
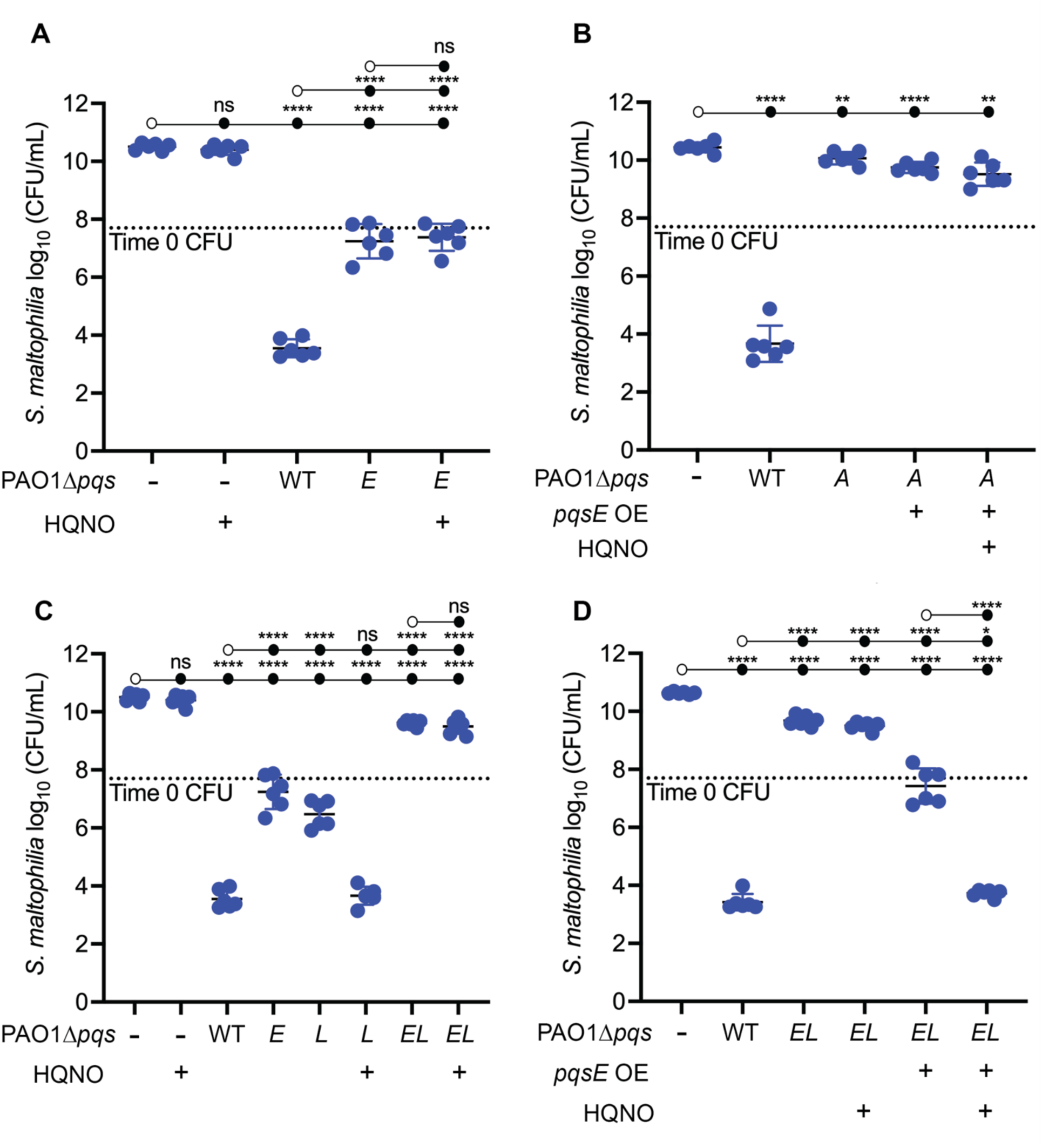
HQNO and PqsE are both required to mediate competition between *P. aeruginosa* and *S. maltophilia*. **A.** Wild-type K279a was grown in mono- or co-culture with wild-type PAO1 or PAO1Δ*pqsE* supplemented with 50 µM HQNO. **B.** Wild-type K279a was grown in mono- or co-culture with wild-type PAO1 or a PAO1Δ*pqsA* mutant overexpressing *pqsE* supplemented with 50 µM HQNO. **C.** Wild-type K279a was grown in mono- or co-culture with wild-type PAO1, PAO1Δ*pqsE*, PAO1Δ*pqsL*, and PAO1Δ*pqsEΔpqsL* supplemented with 50 µM HQNO. **D.** Wild-type K279a was grown in mono- or co-culture with wild-type PAO1, PAO1Δ*pqsEΔpqsL*, and PAO1Δ*pqsEΔpqsL* complemented with *pqsE* supplemented with 50 µM HQNO. All graphs show *S. maltophilia* viability at 24 hours where the x-axis shows each condition and the y-axis shows *S. maltophilia* CFUs. Each experiment was performed in triplicate. Error bars represent the standard deviation. *P* values were calculated between each condition using an unpaired two-tailed t test with Welch’s correction. Statistical comparisons between strains are indicated by a line connecting open and closed circles, with ** denoting *P* < 0.01, *** denoting *P* < 0.001, and **** denoting *P* < 0.0001.

We next asked whether PqsE alone was necessary for HQNO-mediated killing of *S. maltophilia*. To answer this question, we created a *pqsA* knockout mutant that harbored an arabinose-inducible copy of *pqsE* at a neutral location in the chromosome. We competed this mutant against *S. maltophilia* with and without HQNO present (**Figure 5B**). This mutant allowed us to test whether PqsE is important for potentiating HQNO activity in the absence of other alkylquinolones. Surprisingly, we observed no difference in *S. maltophilia* CFUs when it was co-cultured with the *pqsA* mutant or the *pqsA* mutant overexpressing *pqsE*. Further, we observed that the *pqsA* mutant overexpressing *pqsE* did not kill *S. maltophilia* even when HQNO was added. These results indicated that *pqsE* expression is insufficient to mediate competition with *S. maltophilia*, whether or not HQNO is present. These data also are consistent with the idea that some other part of *pqs* QS, in addition to PqsE expression and HQNO production, is necessary for *P. aeruginosa* to kill *S. maltophilia*. Further, *pqs* QS was shown to solely regulate the PQS biosynthesis locus, so it is likely that a product of this locus is necessary for killing *S. maltophilia*.

### HQNO, PqsE, and the PQS biosynthesis pathway are required for mediating competition between *P. aeruginosa* and *S. maltophilia*

We hypothesized that PqsE was required in conjunction with *pqs* QS and HQNO to mediate competition with *S. maltophilia*. To assess this hypothesis, we made a *pqsE* and *pqsL* double knockout mutant (PAO1Δ*pqsEΔpqsL)* that has no PqsE activity or HQNO production; however this mutant can still activate *pqs* QS that includes the production of PQS and intermediate alkylquinolone products (**Figure 5C**). As we observed in earlier experiments, both the *pqsE* and *pqsL* single knockout mutants exhibited partial competitive defects, which could be complemented in the *pqsL* mutant with exogenous HQNO. We observed that the *pqsE* and *pqsL* double knockout mutant had a competitive defect akin to the PQS biosynthesis gene mutants. These data also show that while *pqs* QS is important, it is insufficient and likely requires both PqsE and PqsL for killing *S. maltophilia*.

We further investigated how PqsE and PqsL may be co-implicated in the competion. Although a *pqsL* mutant does not impair PQS and intermediate alkylquinolone production, a *pqsL* mutant has been shown to increase production of the signaling molecules HHQ and PQS [66, 67]. We wondered whether *pqsE* expression was affected in this mutant and performed a quantitative real time PCR experiment to monitor *pqsE* expression in wild-type PAO1, PAO1Δ*pqsA*, PAO1Δ*pqsB*, and PAO1Δ*pqsL* in late-log phase cultures (**Figure 6**). We found that compared to wild-type PAO1, *pqsE* expression decreased by 2-fold in the *pqsA* and *pqsB* deletion mutants. This decrease likely reflects a lack of PqsR activity and therefore reduced induction of the PQS biosynthesis operon. Indeed, the decrease was ameliorated by addition of exogenous PQS to these mutants, showing that the deletions do not dramatically affect mRNA stability (**Supplemental Figure 4**). Unexpectedly, we found that the *pqsL* mutant exhibited a 5-fold increase in *pqsE* expression compared to the wild-type. In the *pqsL* mutant, the enhanced expression of *pqsE* that likely results in increased PqsE activity. This result suggested to us that in the absence of HQNO production, there is likely some competition mediated by the increased *pqs* QS and PqsE activity, but full competition still does require HQNO.

**Figure 6.**
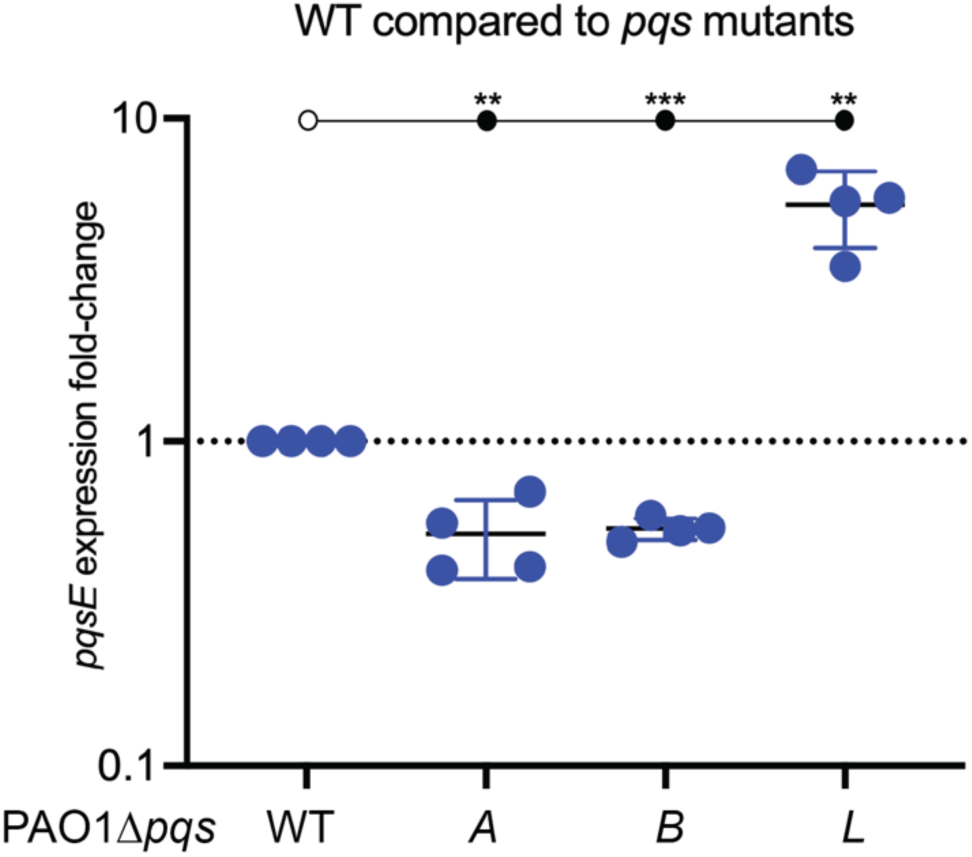
A *pqsL* mutant has increased *pqsE* expression compared to wild-type PAO1. Quantitative real-time PCR measuring *pqsE* transcripts at an OD_600_ of 1.0 in wild-type PAO1, PAO1Δ*pqsA*, PAO1Δ*pqsB*, and PAO1Δ*pqsL* cultures. *pqsE* expression was normalized to *rplU* and reported as 2^-ΔΔCT^. Each experiment was performed in duplicate. Error bars represent the standard deviation. *P* values were calculated between each condition using an unpaired two-tailed t test with Welch’s correction. Statistical comparisons between strains are indicated by a line connecting open and closed circles, with ** denoting *P* < 0.01 and *** denoting *P* < 0.001.

Finally, we tested whether the *pqsE* and *pqsL* mutant could be complemented by the addition of exogenous HQNO (**Figure 5C**). We found that unlike the *pqsL* single knockout mutant, the *pqsE* and *pqsL* double knockout mutant is no longer complemented by the addition of HQNO. These data indicate that a PqsE-regulated factor is required for HQNO-mediated competition with *S. maltophilia*. We further tested whether we could genetically and chemically complement this phenotype (**Figure 5D**). We generated a *pqsE* and *pqsL* double knockout mutant and cloned in a chromosomal copy of *pqsE* (PAO1Δ*pqsEΔpqsL* + *pqsE)* under control of an arabinose-inducible promoter. We relied on the leakiness of the arabinose-inducible promoter in the absence of arabinose addition to allow for *pqsE* expression. We observed some *S. maltophilia* killing when in co-culture with the PAO1Δ*pqsEΔpqsL* + *pqsE* mutant compared to the PAO1Δ*pqsEΔpqsL* mutant. Further, the killing in the PAO1Δ*pqsEΔpqsL* + *pqsE* mutant was similar to that observed in the *pqsL* mutant (**Figure 5C**). In contrast to the PAO1Δ*pqsEΔpqsL* mutant, chemically complementing the PAO1Δ*pqsEΔpqsL* + *pqsE* mutant with exogenous HQNO completely restored the ability of *P. aeruginosa* to kill *S. maltophilia* akin to wild-type PAO1. This experiment shows that *P. aeruginosa* requires both a product made during PQS biosynthesis and a PqsE-regulated gene(s) to enact HQNO-mediated killing on *S. maltophilia*. Taken together, these data demonstrate separate but indispensible roles for each of HQNO, PqsE, and PQS biosynthesis in mediating competitive interactions between *P. aeruginosa* and S. *maltophilia*.

## Discussion

We explored the competition between the Gram-negative bacteria *P. aeruginosa* and *S. maltophilia* and defined a suite of *P. aeruginosa* genes that modulate interactions between these bacteria. Using our co-culture model, we identified that *P. aeruginosa* kills *S. maltophilia*. We showed, using a genetic approach, that this killing depends on *P. aeruginosa* QS. We identified a mechanism where components of the *P. aeruginosa* quinolone QS (*pqs* QS) circuit mediate interactions with another Gram-negative bacterium, a new discovery. We showed that both the alkylquinolone HQNO and the protein PqsE are each necessary but insufficient for competition between the two bacteria. Our data suggest that all three of PQS biosynthesis, HQNO production, and PqsE are required for full killing; removal of any one of these results in significant attenuation (**Figure 7**).

**Figure 7.**
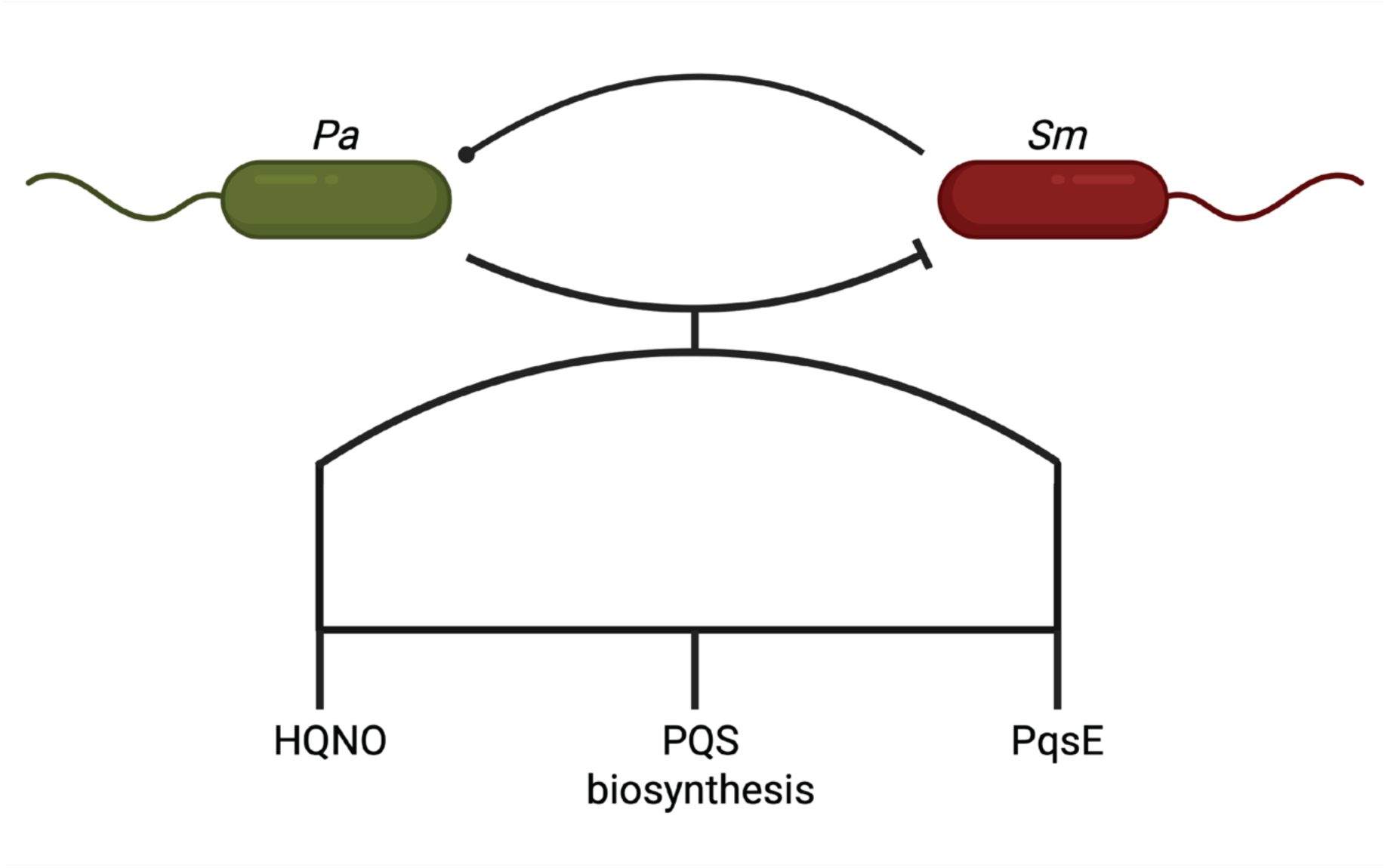
*P. aeruginosa* and *S. maltophilia* interact via competition largely driven by *pqs* QS and PQS biosynthesis. A model showing that the competition in co-culture is driven by an AND logic gate where the *pqs* QS-produced inhibitor HQNO, PQS biosynthesis, and PqsE-mediated activity are required to mediate *S. maltophilia* killing.

Our work offers a new perspective on interactions between *P. aeruginosa* and *S. maltophilia.* Prior studies, in models systems that interrogated growth in a mouse model and in biofilms, established that the interaction between these two bacteria can be cooperative [68, 69]. *S. maltophilia* had increased bacterial load in the presence of viable *P. aeruginosa*; the bacteria together formed integrated biofilms [68]. Further, *S. maltophilia* reduced *P. aeruginosa mobility*, but co-culture induced alginate expression in *P. aeruginosa* and may protect *S. maltophilia* against tobramycin in a mixed biofilm [69]. A recent study found that PQS induced aggregation in *S. maltophilia* grown in a modified minimal medium, and this aggregation conferred protection against *P. aeruginosa* competition [70]. Our experiments established another example that the interactions between these two bacteria can be antagonistic. We showed that *P. aeruginosa* decreases *S. maltophilia* bacterial load through factors, both intrinsic and secreted, regulated by *pqs* QS. These differences highlight the importance of the conditions of experiments on bacterial physiology and interactions; for example, the buffered medium that we use enhances the stability of AHL signals, possibly revealing QS-specific phenotypes.

QS has been shown to regulate several factors involved in interspecies interactions, but our finding of a role for *pqs* QS in competition with a Gram-negative bacterium is new. The RhlR-regulated products hydrogen cyanide, rhamnolipids, and phenazines are shown to mediate competition with other bacteria. Although our data are consistent with a minor role for these factors in mediating competition, they highlight the importance of the alkylquinolone HQNO. While HQNO has been shown to act on other bacteria like *S. aureus*, our work shows a unique mechanism in that HQNO by itself is unable to affect *S. maltophilia* but instead requires other *P. aeruginosa* factors for its activity. HQNO, PqsE-dependent activity, and PQS biosynthesis forms a three-component AND logic gate for *S. maltophilia* killing.

PqsE acts as a chaperone for the transcriptional regulator RhlR and is required for expression of a subset of RhlR-regulated genes [62–65]. For example, RhlR requires PqsE to positively regulate the expression of the phenazine biosynthesis genes, but not the expression of *rhlAB*, genes encoding enzymes that produce rhamnolipids. Our data suggest that in the competition between these species, HQNO activity is potentiated by expression of PqsE, suggesting roles for RhlR-regulated factors (including, possibly, toxins like hydrogen cyanide). Future work may explore PqsE-dependent RhlR-regulated factors that drive competition of *P. aeruginosa* with other Gram-negative bacteria. Another question that arises from our work is what is the other factor driving competition that requires the PQS biosynthesis pathway? A PAO1Δ*pqsA* mutant overexpressing *pqsE* in the presence of HQNO did not kill *S. maltophilia* (**Figure 5B**). This mutant is deficient in the ability to produce a variety of products in the PQS biosynthesis pathway, but we found that the addition of HHQ, HQNO, and PQS are insufficient to restore competition. The branching nature of the PQS biosynthesis pathway makes it difficult to use a genetic approach for the identification of this product or products, but a future determination of the relevant alkylquinolones would likely reveal novel modes of competition mediated by PQS biosynthesis and further our understanding of competitive interactions between bacteria.

It would be interesting to see how these interactions change by varying growth conditions. For example, the role of *pqs* QS in co-culture during biofilm formation is not addressed by our work. Another context would be to replicate the nutritional environment of infection sites, such as by using synthetic cystic fibrosis medium [71]. Beyond the context of infection, experiments could examine the impact of QS, HQNO, PqsE, or PQS biosynthesis in the context of growth in the environment, including in water or the soil, using a variety of environmental isolates.

Our work expands our understanding on how bacteria use QS to mediate interactions, competition, and sociality. We discovered a mechanism for competition between *P. aeruginosa* and *S. maltophilia*, where HQNO acts in concert with PqsE and PQS biosynthesis to kill bacteria. These results have implications that may apply to other interactions. For example, what effect do PqsE and HQNO have on other Gram-negative bacteria? Have Gram-negative bacteria (including *S. maltophilia*) evolved countermeasures to negate the effects of HQNO? Is this quinolone-based mechanism of competition used by bacteria other than *P. aeruginosa*? Our work sets the stage for future studies of how quorum sensing and the environment affect the mechanisms of bacterial competition, and how bacteria respond to these challenges.

## Materials and Methods

### Bacteria and growth conditions

Strains and plasmids used in this study are listed in **Supplemental Tables 1 and 2** *P. aeruginosa* was grown in lysogeny broth (LB) buffered with 50 mM 3-(*N*-morpholino) propanesulfonic acid, pH 7.0 (LB-MOPS). *Escherichia coli* was grown in LB. Cultures were grown in 18-mm test tubes at a volume of 3 mL in a shaking incubator (250 RPM) at 37°C. For individual colony growth we used LB supplemented with 1.5% agar. Where required, broth cultures of *E. coli* and *P. aeruginosa* were supplemented with gentamicin at a concentration of 10 µg per mL (Gm10). *E. coli* colonies were grown on LB supplemented with 1.5% agar and gentamicin at 10 µg per mL. *P. aeruginosa* colonies were grown on LB supplemented with 1.5% agar and gentamicin at 100 µg per mL (Gm100) for transformations.

### Construction of *P. aeruginosa* mutants

In all experiments with the laboratory strain PAO1, we used strain *P. aeruginosa* PAO1-UW [72]. In-frame deletions of *pqsA*, *pqsB*, *pqsC*, *pqsD*, *pqsE*, *pqsH*, and *pqsL* were generated using two-step allelic exchange as previously described [73]. Briefly, constructs for gene deletions were created by using a pEXG2 vector backbone and Gibson assembly to 1000 bp of DNA flanking each side of the gene of interest to facilitate homologous recombination. *E. coli* S17-1 was transformed with each construct and used to deliver knockout plasmids to *P. aeruginosa* via conjugation. Merodiploids were selected by plating on *Pseudomonas* Isolate agar containing Gm100, and deletion mutants were then selected on LB agar containing 15% sucrose and no sodium chloride. All deletion mutants were confirmed by PCR and sequencing of genomic DNA.

Overexpression constructs were created using a pUC18T mini Tn7T integrating plasmid with an arabinose-inducible promoter and Gibson assembly to clone in the *pqsE* CDS downstream of the promoter [74]. The overexpression construct was electrotransformed into *P. aeruginosa* and selected using LB supplemented with 1.5% agar and gentamicin at 100 µg per mL (Gm100) [75]. Proper integration at the attTn7 attachment site was confirmed by PCR and sequencing of genomic DNA. Gentamicin susceptible mutants were made using Flp-mediated excision of gentamicin resistance marker [76].

#### Co-culture assay

*P. aeruginosa* and *S. maltophilia* cultures were inoculated from individual colonies into 18-mm test tubes containing LB-MOPS and grown overnight 37_°_C with shaking. Experiments were performed in biological (different colonies) duplicate. Cultures were back diluted to an OD_600_ of 0.1 and grown until OD_600_ of 0.2. Early exponential phase cultures were used to inoculate 18-mm tubes containing LB-MOPS to an OD_600_ of 0.025 with a 3 mL final volume. Cultures were supplemented with PQS (20 µM), HHQ (50 µM), or HQNO (50 µM) at this time, if required. Cultures were incubated at 37_°_C with shaking for 24 hours. At various time points, 100 µL of culture was sampled and used to measure colony forming units (CFU). To select for *S. maltophilia*, cultures were plated onto LB agar containing 10 µg/mL gentamicin. To select for *P. aeruginosa*, cultures were plated onto cetrimide agar or Columbia agar containing 30 mg/L 9-chloro-9-[4-(diethylamino)phenyl]-9,10-dihydro-10-phenylacridine hydrochloride (C-390) and 30 mg/L phenanthroline [55].

#### RNA isolation

Wild-type PAO1, PAO1Δ*pqsA*, PAO1Δ*pqsB*, and PAO1Δ*pqsL* cultures were started from single colonies in LB-MOPS in biological duplicate. Cultures were incubated for 18 hours at 37_°_C with shaking. Cultures were back diluted to an OD_600_ of 0.025 and grown until cultures reached an OD_600_ of 0.1, and then cultures are diluted back to an OD_600_ of 0.05. At an OD_600_ of 1.0, a total of an OD_600_ of 2.0 was pelleted at 4000 RPM for 5 min. Supernatant was discarded and pellets were resuspended in 1 mL QIAzol containing lysis beads. Samples were lysed by bead-beating at maximum RPM for 1 min with chilling for 5 min. Bead-beating was repeated twice. Chloroform was added, shaken vigorously, and centrifuged for 15 min at 12,000 x g at 4_°_C. 450 µL of the upper phase was combined with 675 µL of 100% ethanol. Samples RNA was extracted using a RNeasy kit with one on-column DNase (cat. No. 79254, Qiagen) treatment, and RNA was eluted using RNase-free water.

#### qRT PCR analysis

RNA was extracted as described above. 250 ng total cDNA was generated by reverse transcription using the qScript cDNA synthesis kit. Quantitative real-time PCR was performed using 2.5 ng total cDNA using the PowerTrack SYBR Green Mix. Primers amplifying a *pqsE* amplicon were used to monitor *pqsE* expression. Primers amplifying a *rplU* amplicon were used as a housekeeping gene (**Supplemental Table 2)**. Relative gene expression levels were determined using the 2^-ΔΔCt^ method.

## Abbreviations

CFU: colony forming unit
CF: cystic fibrosis
Gm: gentamicin
LB: lysogeny broth
PQS: Pseudomonas quinolone signal
QS: quorum sensing
WT: wild-type

## Acknowledgements

We thank Maureen Thomason and Nicole Smalley for their assistance in cloning. This work was funded in part by grants from Cystic Fibrosis Foundation (FRANDO24F0) to AF and the NIH (R35GM152107 and R01AI177575) to AAD.

