## Supplemental figures for "A three-component logic gate governs quorum sensing-regulated killing of *Stenotrophomonas maltophilia* by *Pseudomonas aeruginosa*"

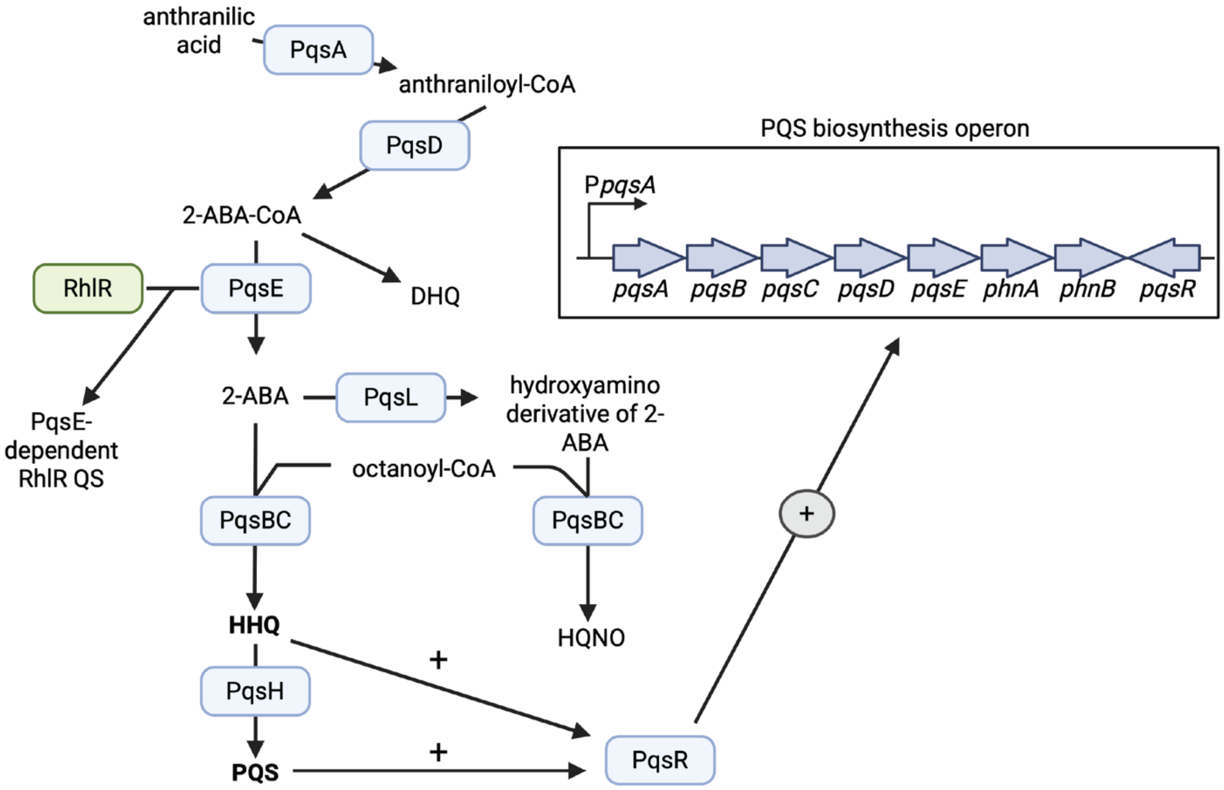


**Supplemental Figure 1. PQS QS and *pqs* biosynthesis pathway.** PQS, HHQ, and intermediates are produced by the activities of the enzymes PqsABCDH. PQS and HHQ bind to and activate the transcriptional regulator MvfR (also known as PqsR). PqsR positively regulates the expression of the *pqs* biosynthesis operon, including *pqsA*, *pqsB*, *pqsC*, *pqsD*, and *pqsE*, and creates a positive feedback loop. Adapted from Giallonardi et al.

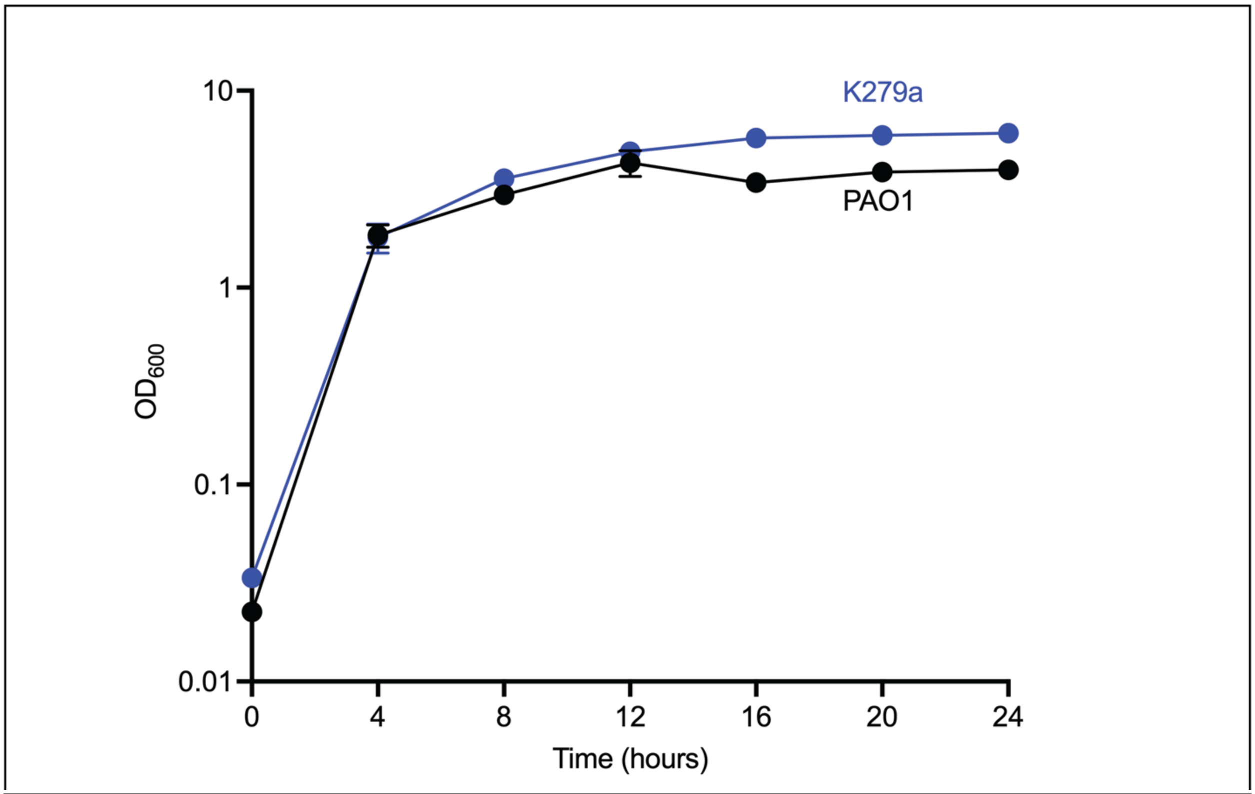


**Supplemental Figure 2.** *P. aeruginosa* and *S. maltophilia* have similar growth rates. Growth of wild-type K279a and PAO1 in buffered LB was measured over 24 hours. The graph shows time in hours on the x-axis and $OD_{600}$ on the y-axis. Each experiment was performed in triplicate. Error bars represent the standard deviation.

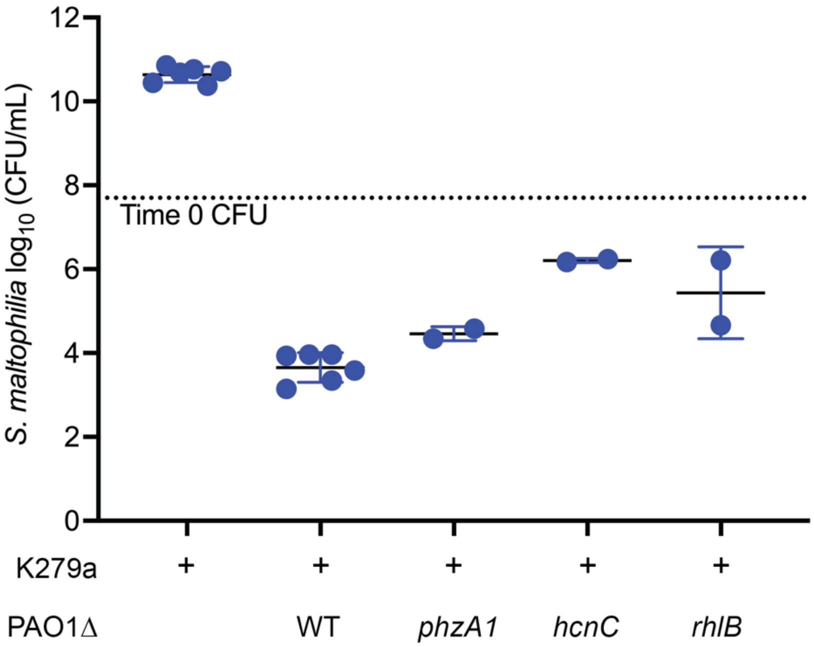


**Supplemental Figure 3. Hydrogen cyanide and rhamnolipids mediate some competition between *Pa* and *Sm*.** Wild-type K279a was grown in mono- or co-culture with wild-type PAO1, PAO1Δ*phzA1*, PAO1Δ*hcnC,* or PAO1ΔrhlB. The graph shows *S. maltophilia* viability at 24 hours where the x-axis shows each condition and the y-axis shows log$_{10}$-transformed *S. maltophilia* CFUs. Each experiment was performed in triplicate. Error bars represent the standard deviation.

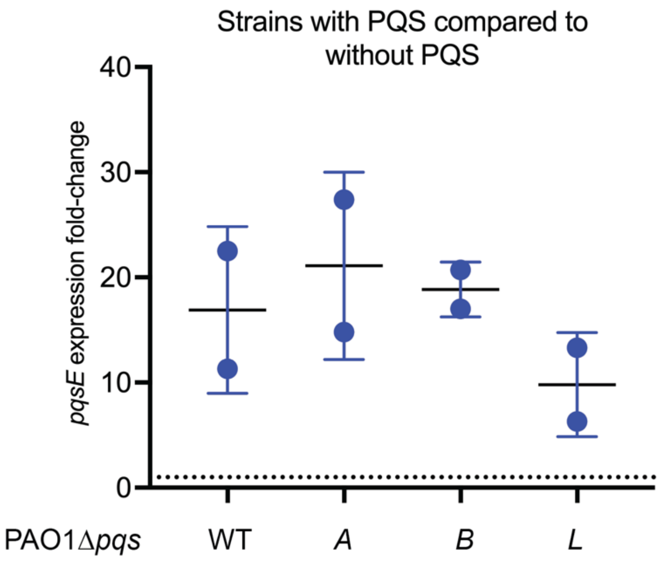


**Supplemental Figure 4. Exogenous addition of PQS induces *pqsE* expression in wild-type and *pqsA, pqsB, and pqsL* knockout mutants.** Quantitative real-time PCR measuring *pqsE* transcripts at an $OD_{600}$ of 1.0 in wild-type PAO1, PAO1Δ*pqsA*, PAO1Δ*pqsB*, and PAO1Δ*pqsL* cultures with and without the addition of 20 µM PQS. For each strain, we compared *pqsE* transcript abundance with PQS addition to cultures without the addition of PQS. *pqsE* expression was normalized to *rplU* and reported as $2^{-\Delta\Delta CT}$.
