## Supplemental tables for "A three-component logic gate governs quorum sensing-regulated killing of *Stenotrophomonas maltophilia* by *Pseudomonas aeruginosa*"

**Supplemental Table 1. Bacterial strains used in this study.**

| Bacterial strain | Description | Source |
|---|---|---|
| ***P. aeruginosa*** | | |
| PAO1 | Wild-type strain | (69) |
| PAO1Δ*lasR* | PAO1 with unmarked, in-frame *lasR* deletion | (74) |
| PAO1Δ*rhlR* | PAO1 with unmarked, in-frame *rhlR* deletion | (75) |
| PAO1Δ*pqsR* | PAO1 with unmarked, in-frame *pqsR* deletion | (75) |
| PAO1Δ*pqsA* | PAO1 with unmarked, in-frame *pqsA* deletion | This work |
| PAO1Δ*pqsB* | PAO1 with unmarked, in-frame *pqsB* deletion | This work |
| PAO1Δ*pqsC* | PAO1 with unmarked, in-frame *pqsC* deletion | This work |
| PAO1Δ*pqsD* | PAO1 with unmarked, in-frame *pqsD* deletion | This work |
| PAO1Δ*pqsE* | PAO1 with unmarked, in-frame *pqsE* deletion | (75) |
| PAO1Δ*pqsH* | PAO1 with unmarked, in-frame *pqsH* deletion | This work |
| PAO1Δ*pqsL* | PAO1 with unmarked, in-frame *pqsL* deletion | This work |
| PAO1Δ*pqsE*Δ*pqsL* | PAO1 with unmarked, in-frame of *pqsE* and *pqsL* deletion | This work |
| ***E. coli*** | | |
| NEB5α | *fhuA2Δ(argF-lacZ)U169 phoA glnV44 Φ80Δ(lacZ)M15 gyrA96 recA1 relA1* | NEB |
| S17-1 | *recA pro hsdR RP4-2Tc::Mu-Km::Tn7* | (76) |

**Supplemental Table 2. Plasmids used in this study.**

| Plasmids | Description | Source |
|---|---|---|
| pEXG2 | Allelic exchange vector with pBR origin, *sacB*, Gm$^{R*}$ | (77) |
| pEXG2-PAO1-*pqsA*-KO | pEXG2 containing sequences for PAO1 *pqsA* in-frame deletion | This work |
| pEXG2-PAO1-*pqsB*-KO | pEXG2 containing sequences for PAO1 *pqsB* in-frame deletion | This work |
| pEXG2-PAO1-*pqsC*-KO | pEXG2 containing sequences for PAO1 *pqsC* in-frame deletion | This work |
| pEXG2-PAO1-*pqsD*-KO | pEXG2 containing sequences for PAO1 *pqsD* in-frame deletion | This work |
| pEXG2-PAO1-*pqsE*-KO | pEXG2 containing sequences for PAO1 *pqsE* in-frame deletion | (75) |
| pEXG2-PAO1-*pqsH*-KO | pEXG2 containing sequences for PAO1 *pqsH* in-frame deletion | This work |
| pEXG2-PAO1-*pqsL*-KO | pEXG2 containing sequences for PAO1 *pqsL* in-frame deletion | This work |

* Gm$^{R}$, resistant to gentamicin
